# Twenty-one Year Trends for Shorebirds, Waterfowl, and Other Waterbirds at Great Salt Lake, Utah

**DOI:** 10.1101/2021.05.17.444474

**Authors:** Brian G. Tavernia, Tim Meehan, John Neill, John Luft

## Abstract

Millions of wetland-dependent birds annually depend on saline lakes and associated wetlands in the western United States. Understanding the population status and trends of birds with different life histories and habitats can guide efforts to secure water resources needed to sustain bird habitats. We used a 21-year dataset to examine population trends for 24 survey units presumed to be high-quality habitat for migratory shorebirds, waterfowl, and other waterbirds at Great Salt Lake and associated wetlands. As expected for high-quality habitats, we found stable or positive trends for 36 of 37 species or groups in fall, spring, or both seasons when considering survey units in aggregate. Despite stable or positive aggregate trends, negative trends did occur in some individual survey units. Foraging, migration distance, and taxonomic groupings were unrelated to trend direction. Research is needed to test whether survey units represent high-quality habitat. With declining regional water resources, stable and positive aggregate trends reinforce the importance of surveyed units at Great Salt Lake and associated wetlands to wetland-dependent birds. Ensuring continuation of stable and positive trends will require identifying environmental factors - including water quantity and quality - driving trends, and require coordinated regional management and monitoring of wetland-dependent birds.

## Introduction

The saline lakes of the western United States and their associated wetlands support millions of shorebirds, waterfowl, and other waterbirds on an annual basis (Aldrich and Paul 2002; Petrie *et al*. 2013; Wilsey *et al*. 2017). Bird use of these systems is driven by multiple factors including the predominantly xeric conditions of the western United States, spatially and temporally dynamic water depths, diverse salinities and dynamic wetland habitats, the presence of islands for nesting within lakes, and abundant food resources (Aldrich and Paul 2002; Wilsey *et al*. 2017; Sorensen *et al*. 2020). The factors that determine habitat value are affected indirectly or directly by saline lakes’ water levels, which depend on the balance among water inflows, precipitation, and evaporative water loss. For example, brine shrimp (*Artemia* spp.), an important invertebrate food resource at some lakes, are sensitive to salinity changes caused by receding or rising lake levels (Dana and Lenz 1986; Senner *et al*. 2018). Water diversions and extractions for anthropogenic uses, such as irrigated agriculture, have historically reduced water inflows to saline lakes and associated wetlands (Wurtsbaugh *et al*. 2017; Donnelly *et al*. 2020) and, in combination with climate change (e.g., potential for reduced streamflow), will continue to affect water levels, timing, and salinity in the future (Ficklin *et al*. 2013; Jeppesen *et al*. 2015; Meixner *et al*. 2016). Efforts to protect and restore water flows for saline lakes and associated wetland habitat can benefit from understanding the population status and trends of bird species with different life histories and habitat requirements.

Population trend assessments depend on monitoring, or the process of making and analyzing repeated observations of species’ attributes to track changes in their status across time (Thompson *et al*. 1998). By supplying estimates of species distributions, population sizes, or trends, monitoring data inform species prioritization for conservation actions given available resources. Multiple prioritization systems assign greater priority to species with smaller distributions and population sizes and declining trends, and conservation plans have used these factors to prioritize shorebird and waterbird species (Brown *et al*. 2001; Kushlan *et al*. 2002). Beyond species prioritization, trend assessments can be used to suggest conservation strategies and tactics to benefit species and the habitats on which they rely. The potential for these suggestions is realized if researchers identify life history (e.g., migration strategy) or ecological (e.g., habitat use) traits shared in common by species with decreasing, stable, or increasing trends. For example, wetland birds capable of using artificial waterbodies, such as impoundments, may increase with agricultural and urban development in arid regions whereas those dependent on natural wetlands may decrease (Okes *et al*. 2008). In such regions, conservationists may focus on protecting remaining natural wetlands and restoring degraded areas by, for example, managing water inflows.

Trends for specific sites may not reflect the overall trajectory of a species’ regional population if surveyed sites are not representative of available habitat, and this must be weighed when interpreting trends to inform conservation and management actions. As one example, populations have been shown to be relatively invariable in perceived high-quality habitat when compared to population fluctuations in lower quality sites (Kluyver and Tinbergen 1953; Gill *et al*. 2001). Lower quality sites are said to ‘buffer’ fluctuations in higher quality sites, and this buffering could result from differences in survival, reproduction, or active habitat selection of high-quality sites (Kluyver and Tinbergen 1953). Thus, lower quality sites potentially reflect changes in overall population size to a greater degree than higher quality areas (Gill *et al*. 2001). Accordingly, trends based on surveys of high-quality habitat sites may not detect overall population declines until such declines have progressed enough to become apparent in high-quality habitat. Similarly, omission of some high-quality habitat types (e.g., shorebird playa habitat), such that sampled habitat types are incomplete, may result in survey trends that do not reflect trends in the regional population.

Great Salt Lake, the largest saline lake in the Great Basin, and its associated wetlands are recognized regionally, nationally, and hemispherically as important sites for shorebirds, waterfowl, and other waterbirds (Aldrich and Paul 2002; Chipley *et al*. 2003). As with other saline lakes, anthropogenic water diversions have reduced water inflows to Great Salt Lake and associated wetlands (Wurtsbaugh *et al*. 2017), and climate change may cause regional shifts from snow to rainfall, changes in snowmelt timing, and increased evapotranspiration that contribute to water inflow reductions in the future (Baxter and Butler 2020). Local and regional plans and other documents have identified priority migratory shorebird, waterfowl, and other waterbird species that depend on Great Salt Lake and have called for quantifying flows necessary to provide habitat for these species (Ivey and Herziger 2006; Petrie *et al*. 2013; Sorensen *et al*. 2018).

In this study, we used a 21-year monitoring dataset to analyze trends of migratory shorebird, waterfowl, and other waterbirds in high bird-use areas of Great Salt Lake and its associated wetlands. In conducting analyses, our objectives were to (1) estimate population trends during fall and spring migration for individual species and species groups; (2) determine whether trends differed for different areas of the lake and associated wetlands; and (3) evaluate whether trends were associated with particular taxonomic groups, migratory strategies, or foraging techniques. With a focus on high-use areas of presumably high habitat quality, we predicted most species and species groups would show stable trends. Ultimately, we aimed to provide trend estimates that can inform future discussions about species prioritization and to identify traits shared by species showing increasing, decreasing, or stable trends to inform the formulation of conservation strategies.

## Methods

### Study Area

Unless another citation is provided, the description of Great Salt Lake (Fig. 1) is based on information in Aldrich and Paul (2002) who described the lake’s ecological setting from an avian perspective. The lake is located in a cold desert environment with local annual precipitation ranging from 25 cm to 38 cm on its west and east sides, respectively. Temperatures frequently reach −18°C during winter and 38°C during summer. The lake is one of a relatively few locations where migratory wetland birds may find habitat for staging and molting in an otherwise arid region. As with other saline lakes, water enters through surface water, groundwater flows, and precipitation and naturally exits via evaporation (Null and Wurtsbaugh 2020). These hydrological processes have led and continue to lead to the accumulation of salts in the lake. Freshwater inflows predominantly from the Bear, Weber, and Jordan rivers into this terminal lake create a continuum of freshwater, brackish, and saline wetland habitats for bird use. Other inflows are from smaller tributaries, groundwater, sewage plants, and precipitation. Based on National Wetlands Inventory data (U.S. Fish and Wildlife Service 2019), the 5 counties (Box Elder, Davis, Salt Lake, Tooele, and Weber) spatially adjacent to and including Great Salt Lake contain 1.4 million ha of lake; 70,583 ha of freshwater emergent wetland; 22,335 ha of riverine wetlands; 8,357 ha of freshwater pond; and 1,056 ha of freshwater forest/shrub wetland.

**Figure 1.**
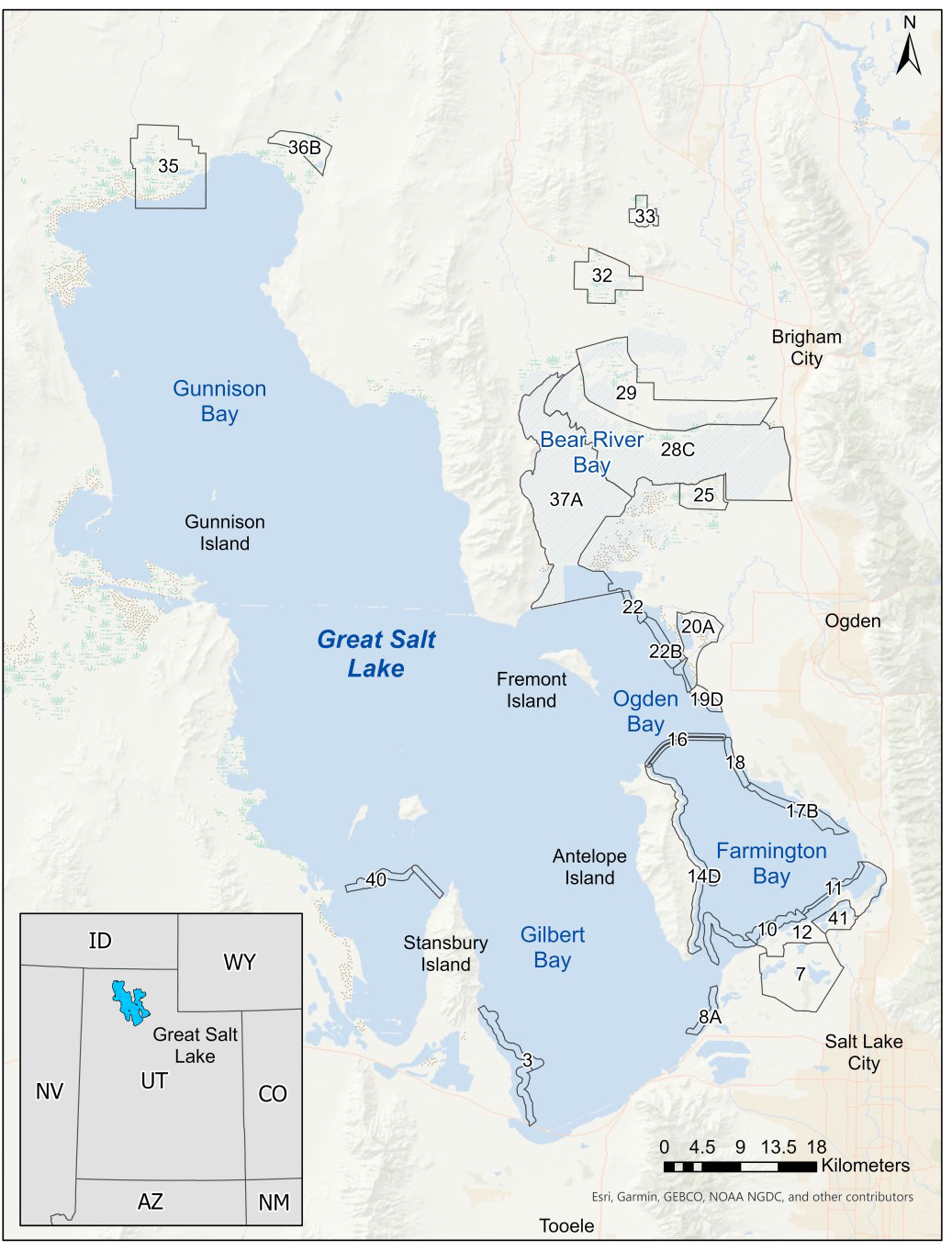
Locations of 24 survey units where the Great Salt Lake Ecosystem Program conducts long-term surveys for migratory shorebirds, waterfowl, and other waterbirds at Great Salt Lake, Utah

The plant composition of Great Salt Lake wetlands responds to salinity, hydroperiod, and water depth. Freshwater wetlands tend to host plant species such as cattail (*Typha* spp.), sago pondweed (*Stuckenia pectinata*), and hardstem bulrush (*Schoenoplectus acutus*) whereas examples from more saline wetlands include muskgrass (*Chara* spp.) and alkali bulrush (*Bolboschoenus maritimus*). Shorebirds will use mudflats vegetated by pickleweed (*Salicornia* spp.) and wet meadows with saltgrass (*Distichlis spicata*) (Sorensen *et al*. 2020). Invasion by common reed (*Phragmites australis*) can reduce or eliminate wetland habitat suitability for birds (Benoit and Askins 1999), and managers expend considerable effort and great expense to control such invasions at Great Salt Lake (Rohal *et al*. 2018). Shorebird, waterfowl, and other waterbird use of wetland plants and habitats more generally at Great Salt Lake is reviewed by Sorensen *et al*. (2020) and Downard et al. (2017). Bird diets at Great Salt Lake may be primarily herbivorous, insectivorous, piscivorous, or omnivorous with dietary status varying from species-to-species and season-to-season (Barber and Cavitt 2012). Some birds, such as the Eared Grebe (*Podiceps nigricollis*) and Wilson’s Phalarope (*Phalaropus tricolor*), are particularly dependent on halophilic brine shrimp (*A. franciscana*) and brine flies (*Ephydra* spp.) as food resources (Conover and Caudell 2009; Roberts 2013).

Variable inflows, evaporative water loss, lake surface elevation, and a low gradient bottom interact and result in changes in the types and extents of wetland habitats present over seasonal, annual, and decadal periods. As an example, an increase in lake elevation from 1,277.5 m (observed in 1963) to 1,283.8 m (1986) more than doubles the surface area of the lake (Cruff 1986). These water dynamics generally create productive ecological conditions for migratory birds, but extreme fluctuations can negatively affect birds for a short time by, for example, flooding nesting areas. These fluctuations are overlaid on a downward trend in lake level. Since the arrival of European settlers in 1847, water development and river diversions for agricultural, industrial, and urban purposes have reduced water flow into the lake, resulting in the lake being approximately 3.4 m lower in elevation than it otherwise would have been (Wurtsbaugh *et al*. 2017). Continued anthropogenic water use and climate change have the potential to drive additional declines in lake elevation in the future. Recent projections suggest that precipitation and temperature changes are capable of relatively large impacts on lake elevation whereas water conservation efforts (e.g., increased municipal water and industrial use efficiency) can have a positive, although relatively small, effect on lake elevation (Jacobs Engineering Group Inc. 2019).

### Bird Survey Data

Observers counted birds at 24 survey units within Great Salt Lake or wetlands associated with the lake (Fig. 1, Table 1). We limited surveys to bird species in the following families: *Gaviidae, Podicipedidae, Pelecanidae, Phalacrocoracidae, Ardeidae, Threskiornithidae, Anatidae, Rallidae, Gruidae, Charadriidae, Recurvirostridae, Scolopacidae*, and *Laridae*. We selected survey unit locations representing dike edge, open water, shoreline, and wetland habitats in areas heavily used by species in these families (Paul and Manning 2002). We established survey unit boundaries (median size: 1,495.8 ha; Table 1) based on the edges of habitat patches and the ability to complete surveys within 4 hours.

**Table 1.** Characteristics of 24 survey units where the Great Salt Lake Ecosystem Program conducts long-term surveys for migratory shorebirds, waterfowl, and other waterbirds at Great Salt Lake, Utah. Habitat indicates whether a unit is predominantly dike edge (D), open water (O), shoreline (S), or wetland (W). The area of each unit is reported with the percentage of the unit surveyed reported parenthetically. The years in which the unit was surveyed and included in trends analyses are reported.

| Unit | Habitat | Area (ha) | Survey Years Analyzed |
| --- | --- | --- | --- |
| 3 | S | 1,725.8<br>(100) | 1997 – 2001; 2004 – 2017 |
| 7 | W | 5,910.5<br>(15) | 1997 – 2001; 2004 – 2017 |
| 8A | S | 580.3<br>(100) | 1997 – 2001; 2004 – 2006; 2009; 2012; 2015 |
| 10 | W | 768.1<br>(100) | 1997 – 2001; 2004 – 2017 |
| 11 | W/D | 522.2<br>(100) | 1997 – 2001; 2004 – 2007; 2010; 2013; 2014; 2016 |
| 12 | W | 4,544.5<br>(65) | 1997 – 2001; 2004 – 2006; 2008 – 2017 |
| 14D | S | 3,509.5<br>(100) | 1997 – 2001; 2006 – 2017 |
| 16 | D/S | 849.8<br>(100) | 2004 – 2017 |
| 17B | S | 1,331.8<br>(100) | 1997 – 2001; 2004 – 2006; 2009; 2012; 2015 |
| 18 | S | 525.9<br>(100) | 1997 – 2001; 2004 – 2006 |
| 19D | W | 569.3<br>(85) | 2004 – 2017 |
| 20A | W | 2,495.6<br>(60) | 2004 – 2017 |
| 22 | S | 389.4<br>(100) | 1998 – 2001; 2007; 2010; 2013; 2016 |
| 22B | W/S | 850.5<br>(100) | 2007; 2010; 2013; 2016 |
| 25 | W | 1,773.4<br>(33) | 1997 – 2001; 2004 – 2012; 2014 – 2017 |
| 28C | W | 19,870.9<br>(100) | 2004 – 2017 |
| 29 | W | 10,449.4<br>(30) | 1997 – 1999; 2004 – 2007; 2009 – 2017 |
| 32 | W | 3,248.7<br>(20) | 1997 – 2001; 2004 – 2006; 2008; 2011; 2014; 2017 |
| 33 | W | 863.4<br>(35) | 1997 – 2001; 2008; 2011; 2014; 2016 – 2017 |
| 35 | W | 7,607.9<br>(4) | 1997; 2001; 2004 – 2007; 2010; 2013 – 2014; 2016 |
| 36B | W | 1,659.8<br>(40) | 1998 – 2001; 2004 – 2006; 2009; 2012; 2015 |
| 37A | O | 17,955.2<br>(100) | 2004 – 2017 |
| 40 | D | 752<br>(100) | 1998 – 2001; 2004 – 2006 |
| 41 | W | 1,200.2<br>(50) | 1999; 2001; 2004 – 2006; 2009; 2012; 2015 |

Depending on habitat and means of access, observers used either area counts or aerial surveys to count birds within survey units (Table 1). For some units, area or aerial counts were conducted for separate subunits and these counts later aggregated into a total count for the unit (see below). Area counts involved recording all birds seen or heard within a unit while traveling standardized routes (e.g., dike roadways) or transects. Area counts might not cover the entire survey unit due to inaccessibility or visual obstruction by emergent vegetation. The number of observers for an area count depended on challenges associated with the unit, such as unit size and the number of birds typically present. For area counts of shoreline habitats, observers traveled transects. Transects began at a designated starting point and paralleled the shoreline at a perpendicular distance of approximately 91 m. Observers recorded all birds within a 402-m buffer while traveling transects. Across surveys within and between years, observers shifted transects perpendicularly to maintain a distance of 91 m from the shoreline as Great Salt Lake’s water level waxed and waned. From 1997 – 2001, observers stopped at randomly selected points along shoreline transects to conduct 10-minute point counts. This practice was discontinued in following years.

Observers aerially surveyed two large units within Bear River Bay. For aerial surveys, we established 463-m wide transects spaced approximately 1,852-m apart within survey units. During flights, observers used a GPS unit to locate transect start and ending points. Flight speeds and altitudes ranged from 129 to 161 km/h and 24 to 61 m, respectively. Counts were conducted by two observers with each observer counting species observed out to 231.5 m from the plane. Since transects in aggregate covered only a quarter of the survey unit, we extrapolated transect counts to the entire survey unit by multiplying total counts across transects by a factor of four.

Observers included professional wildlife biologists, field technicians, and volunteers. Across years, 17 of 24 survey units were consistently counted by either wildlife biologists and their field technicians or volunteers whereas professional status was inconsistent for the other survey units. While individual observers changed with time, sometimes annually, wildlife biologists or volunteers with long-standing experience generally supervised counts for their respective units each year. We assumed that changes in professional status and observer identity would add noise to data but not result in biased trends.

Surveys varied in frequency and duration across three phases. From 1997 through 2001, observers annually surveyed units every 10 days. In 1997, surveys began in late June and lasted through mid-September whereas, for the remaining years, surveys began in April and lasted through September (for 17 total surveys). Between 2004 and 2006, annual surveys occurred three and nine times during spring (15 April – 14 May) and fall (8 July – 5 September) migration seasons, respectively. Since 2007, surveys have been conducted twice during spring (10 April – 9 May) migration and three times during fall migration (18 July – 31 Aug) with some units surveyed on an annual basis and others triennially. Not all survey units were surveyed during each of the three phases or with the same frequency (Table 1), so the number of years with survey data ranged from 4 to 19 (median: 13.5 years) across survey units.

To enable trends analyses, we identified and retained surveys conducted only during periods common to all three phases. There were five such periods, including Period 1, from 10 April through 24 April; Period 2, from 25 April through 9 May; Period 3, from 18 July through 1 August; Period 4, from 2 August through 16 August; and Period 5, from 17 August through 31 August. Periods 1 and 2 were classified as spring migration counts, and Periods 3, 4, and 5 were classified as fall migration counts. When multiple counts occurred for a survey unit during a survey period, we averaged counts and rounded to the nearest integer value. For those units with surveys conducted in subunits, we averaged subunit counts per period and then summed the averaged subunit counts. After binning counts into the five common survey periods, we aggregated counts a final time into the two migration seasons by taking the maximum count from periods 1 and 2 for spring migration and the maximum count from periods 3, 4, and 5 for fall migration.

We eliminated 77 species or groups that were rare (i.e., observed on < 20 percent of counts) because modeling of sparse data was not expected to yield robust trend estimates. The list of eliminated species and groups is available upon request. We aggregated counts for 20 species we considered difficult for observers to distinguish in the field, producing counts for seven species groups (Table 2). For example, Clark’s Grebe (*Aechmophorus clarkii*), Western Grebes (*A. occidentalis*), and Clark’s/Western Grebe were grouped together and labeled *Aechmophorus*. Counts for 30 other species remained disaggregated during trends analyses (Table 3). Thus, we retained 37 species or groups, and, for each species or group, the maximum count per survey unit, migration season, and year was used for trend analysis.

**Table 2.**
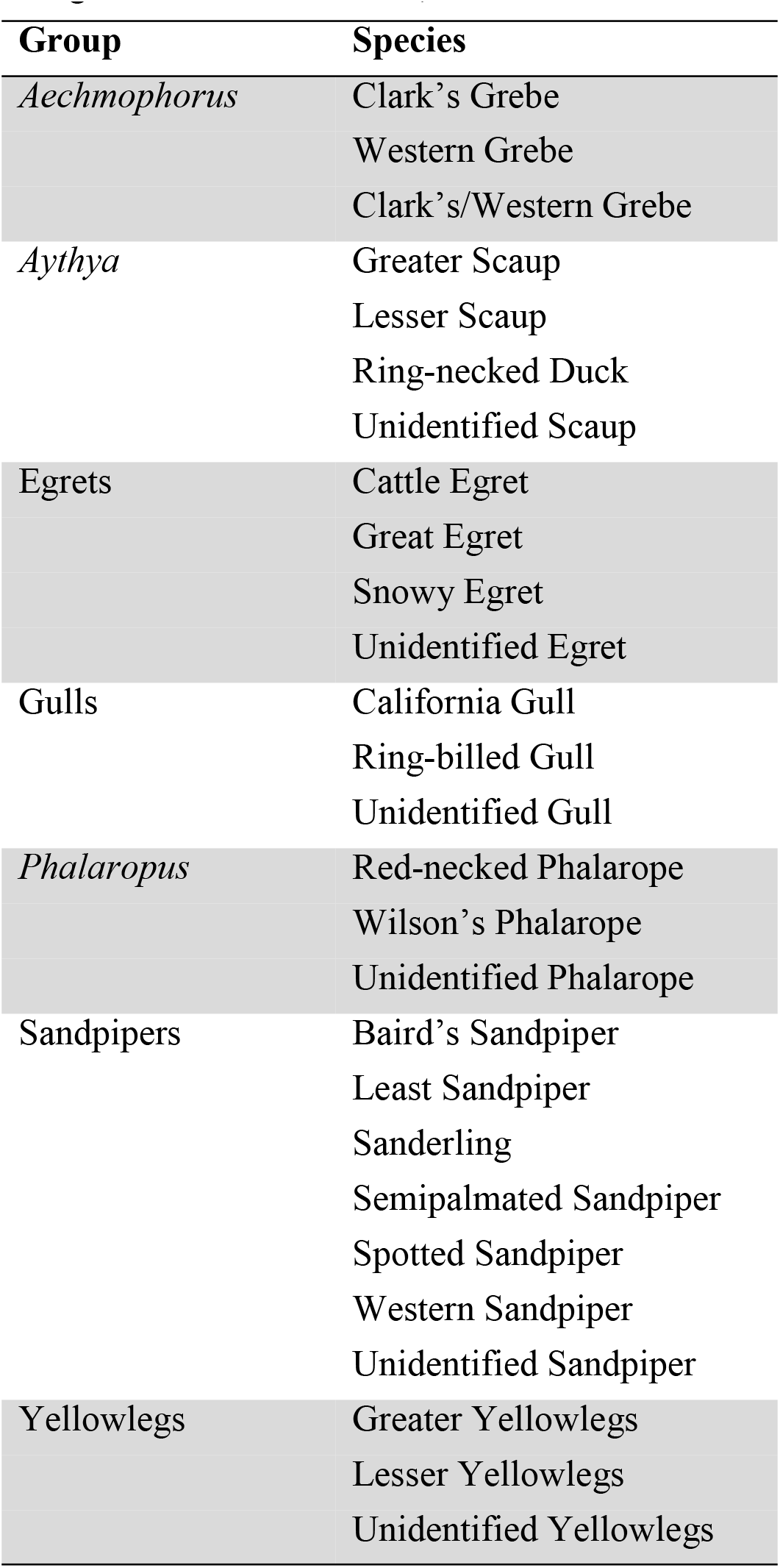
Groups used to aggregate counts for species difficult to identify in the field during surveys conducted by the Great Salt Lake Ecosystem Program at Great Salt Lake, Utah.

**Table 3.** Species-specific foraging techniques, migration distances, and taxonomic groups. Foraging techniques after Ehrlich et al. (1988) and modified based on Cavitt (2006), Barber and Cavitt (2012), Roberts (2013), and Rodewald (2015). Migration distance represents the distance between centroids of nonbreeding and breeding ranges within North and South America as represented by Birdlife International and Handbook of the Birds of the World (2018). For Black-necked Stilt, nonbreeding and breeding ranges were digitized from Cornell Lab of Ornithology (2020). Species migrating ≤ 2000 km were considered short-distance migrants whereas those migrating > 2000 km were labeled long-distance migrants. Taxonomic group assigned based on Clements et al. (2019). Shorebird included species from the order Charadriiformes with the exception of family Laridae whereas waterfowl encompassed species within the order Anseriformes. All other species were considered other waterbirds.

| Species | Foraging Technique | Migration Distance (km) | Taxonomic Group |
| --- | --- | --- | --- |
| American Avocet | Sweeps | 1,821 | Shorebird |
| American Coot | Surface Dips | 994 | Waterbird |
| American Wigeon | Dabbles | 2,796 | Waterfowl |
| American White Pelican | Surface Dips | 2,468 | Waterbird |
| Black-crowned Night-Heron | Stalk and Strike | 1,605 | Waterbird |
| Black-necked Stilt | Ground Glean | 1,319 | Shorebird |
| Canada Goose | Surface Dips | 1,544 | Waterfowl |
| Caspian Tern | High Dives | 3,001 | Waterbird |
| Cinnamon Teal | Surface Dips | 2,178 | Waterfowl |
| Double-crested Cormorant | Surface Dives | 1,228 | Waterbird |
| Eared Grebe | Surface Dives | 1,500 | Waterbird |
| Forster's Tern | High Dives | 2,327 | Waterbird |
| Franklin's Gull | Ground Glean | 9,028 | Waterbird |
| Gadwall | Dabbles | 1,246 | Waterfowl |
| Great Blue Heron | Stalk and Strike | 1,218 | Waterbird |
| Green-winged Teal | Dabbles | 2,442 | Waterfowl |
| Killdeer | Ground Glean | 1,437 | Shorebird |
| Long-billed Curlew | Ground Glean | 1,937 | Shorebird |
| Long-billed Dowitcher | Probes | 5,640 | Shorebird |
| Marbled Godwit | Probes | 2,290 | Shorebird |
| Mallard | Dabbles | 1,521 | Waterfowl |
| Northern Pintail | Dabbles | 3,392 | Waterfowl |
| Northern Shoveler | Surface Dips | 2,949 | Waterfowl |
| Pied-billed Grebe | Surface Dives | 1,256 | Waterbird |
| Redhead | Surface Dives | 1,958 | Waterfowl |
| Ruddy Duck | Surface Dives | 1,169 | Waterfowl |
| Sandhill Crane | Ground Glean | 3,299 | Waterbird |
| Snowy Plover | Ground Glean | 1,468 | Shorebird |
| White-faced Ibis | Ground Glean | 797 | Waterbird |
| Willet | Ground Glean | 3,684 | Shorebird |

### Species Traits

Previous researchers have used nesting, foraging, and migration behaviors to evaluate the response of wetland bird species and communities to natural or anthropogenic disturbances (DeLuca *et al*. 2004; Crewe and Timmermans 2005). With a focus on migration seasons, we categorized species based on their foraging techniques, migration strategy, and taxonomic group (Table 3). We did not categorize species groups because foraging technique and migration strategy varied across a group’s constituents in some cases. We assigned a primary foraging technique to each species after Ehrlich *et al*. (1988) who made assignments based on the breeding season. As our focus was on birds during migration, we revised assigned techniques as needed based on nonbreeding foraging behaviors and food items reported in species accounts of Rodewald (2015) and by the studies of Cavitt (2006), Barber and Cavitt (2012), and Roberts (2013).

We identified species as being either short- (≤2000 km) or long-distance (>2000 km) migrants (Zaifman *et al*. 2017). We followed a procedure for determining migration distance similar to Galbraith *et al*. (2014). Specifically, we quantified the distance between centroids of nonbreeding and breeding ranges as represented by geospatial data of Birdlife International and Handbook of the Birds of the World (2018). Nonbreeding ranges included areas where species were known or thought to be extant, native, and present throughout the year or during the nonbreeding season. Breeding ranges included areas where species were known or thought to be extant, native, and present throughout the year or during the breeding season. We clipped ranges to the boundaries of the North and South American continents (ESRI 2019). The Black-necked Stilt (*Himantopus mexicanus*) was not included in Birdlife International and Handbook of the Birds of the World (2018). We manually digitized this species’ geographic range from North and South America using maps made available by Cornell Lab of Ornithology (2020). All geospatial processing and analyses were carried out in ArcGIS Pro 2.4.1 (ESRI, Redlands, CA).

We assigned species to broad taxonomic categories of shorebird, waterfowl, and other waterbirds based on Clements *et al*. (2019). The shorebird category included species from the order Charadriiformes with the exception of family Laridae whereas waterfowl encompassed all species within the order Anseriformes. All other species were considered waterbirds (Table 3).

### Statistical Analysis

For each species and group, we conducted two independent trend analyses, one using maximum counts from the spring migration season, and one using maximum counts from the fall migration season. We modeled each maximum count, *y_i,t_*, per survey unit *i* during year *t*, as a random variable from a negative binomial distribution. Expected values for maximum counts per study area and year, *μ_i,t_*, were modeled with the linear predictor, log(*μ_i,t_*) = *β_0_* + *β*_1_*Y* + *υ_t_* + *κ_i_* + *τ_i_*, where *β_0_* was a global intercept, *β_1_* was a global log-linear effect of year *Y, υ_t_* was a random intercept deviation per year *t, κ_i_* was a random intercept deviation per study area *i*, and *τ_i_* was a random slope deviation per study area *i*.

The model was analyzed in a Bayesian context using the *INLA* v20.03.17 package (Rue *et al*. 2017) for R statistical computing software (R Core Team 2020). The two global effects were assigned normal prior distributions with a mean = 0 and SD = 100. The three random terms were unstructured, zero-centered, normally distributed, exchangeable effects with penalized complexity priors for the spreads of distributions (Simpson *et al*. 2017). The penalized complexity priors for random intercept deviations were specified such that probability of an SD ≥ 2.00 was 0.01. The penalized complexity prior for random slope deviations was specified such that probability of an SD ≥ 0.50 was 0.01. A negative binomial distribution was adopted for counts to accommodate overdispersion relative to Poisson distributions. Model fits were evaluated by inspecting conditional predictive ordinate distributions for uniformity (Czado *et al*. 2009) and calculating simple correlations between observed and predicted counts per site, year, and season. Models fit reasonably well with correlation coefficients averaging 0.78 (SD = 0.12).

*β_1_* represented the average year effect across all survey units. It became apparent over the course of the analysis that there was considerable variation in temporal trends across survey units, and it was common to find trends for a species or group with positive, negative, and stable trends when examining survey units within which birds varied dramatically in relative abundance. To produce trend estimates weighted by species’ or groups’ relative abundances in survey units, we computed a composite year effect. To calculate a composite year effect, we sampled posteriors of model parameters (n = 10,000) and used the linear predictor to calculate a relative abundance per survey unit and year per sample. Then, for each sample, we summed the relative abundances per year across all survey units, and regressed that estimate against year. This produced the trend estimate reported for the aggregate of all survey units, a strategy common in trend analyses of data from other community science programs (Sauer and Link 2011; Soykan *et al*. 2016). In addition to this aggregate trend, we also computed local effects of year per survey unit using the linear predictor and posterior samples from global and random effects. Local abundance indices were computed as the sum of the global intercept and local intercept deviations. Local year effects were computed as the sum of the global year effect and local effect deviations.

We used two-way contingency tables to evaluate associations between fall and spring aggregate trend directions and species’ foraging techniques, migration strategies, and taxonomic groups. With respect to trend direction, we classified aggregate trend data into two categories, positive or no trend. Only the Redhead (*Aythya americana*) had a negative aggregate trend observed in fall (see Results). Consequently, we did not include a negative direction category, and we dropped the single negative trend from our contingency table analyses. To increase numbers of species associated with foraging techniques, we lumped all species employing diving into a single category, aggregated species using surface dips and dabbles, and combined ground gleaners and probers. We dropped the sweeps and stalk and strike categories due to small numbers of species and the lack of a clear case for aggregation with other foraging techniques. Because ≥ 20% of categories for foraging technique and migration strategy contingency tables had expected values < 5, we did not test the significance of associations using asymptotic χ^2^ distributions (Quinn and Keough 2002). Instead, we compared χ^2^ statistics to χ^2^ distributions generated via Monte Carlo resampling (10,000 resamples) of observed data as implemented in the R package *coin* (Hothorn *et al*. 2006; Hothorn *et al*. 2008).

## Results

### Fall

Of 37 species or groups, four displayed significant positive aggregate population trends during fall with posterior distribution median estimates ranging from 5.87 to 13.54% (Fig. 2, Table 4). One species, the Redhead, had a significant negative population trend of −7.41% during fall. All remaining species or groups had stable aggregate fall trends. When looking at local trends for individual survey units, 17 species or groups possessed significant positive fall trends in up to five survey units, and these 17 included all four species or groups showing significant positive aggregate trends (Fig. 2, Table 4). The survey unit along the east side of Antelope Island had the greatest number of species or groups with positive local trends (Fig. 3). Negative local fall trends occurred in up to 10 survey units for 20 species or groups, and of these 20, one possessed a positive aggregate trend (Table 4). Farmington and Ogden bays held several survey units with large numbers of negative local fall trends for species and species groups (Fig. 3). There were no associations between trend direction and foraging technique (χ^2^_2,26_ = 0.20, p = 1.00), migration strategy (χ^2^_1,29_ = 2.72, p = 0.23), or taxonomic group (χ^2^_2,29_ = 1.60, p = 0.59).

**Fig. 2.**
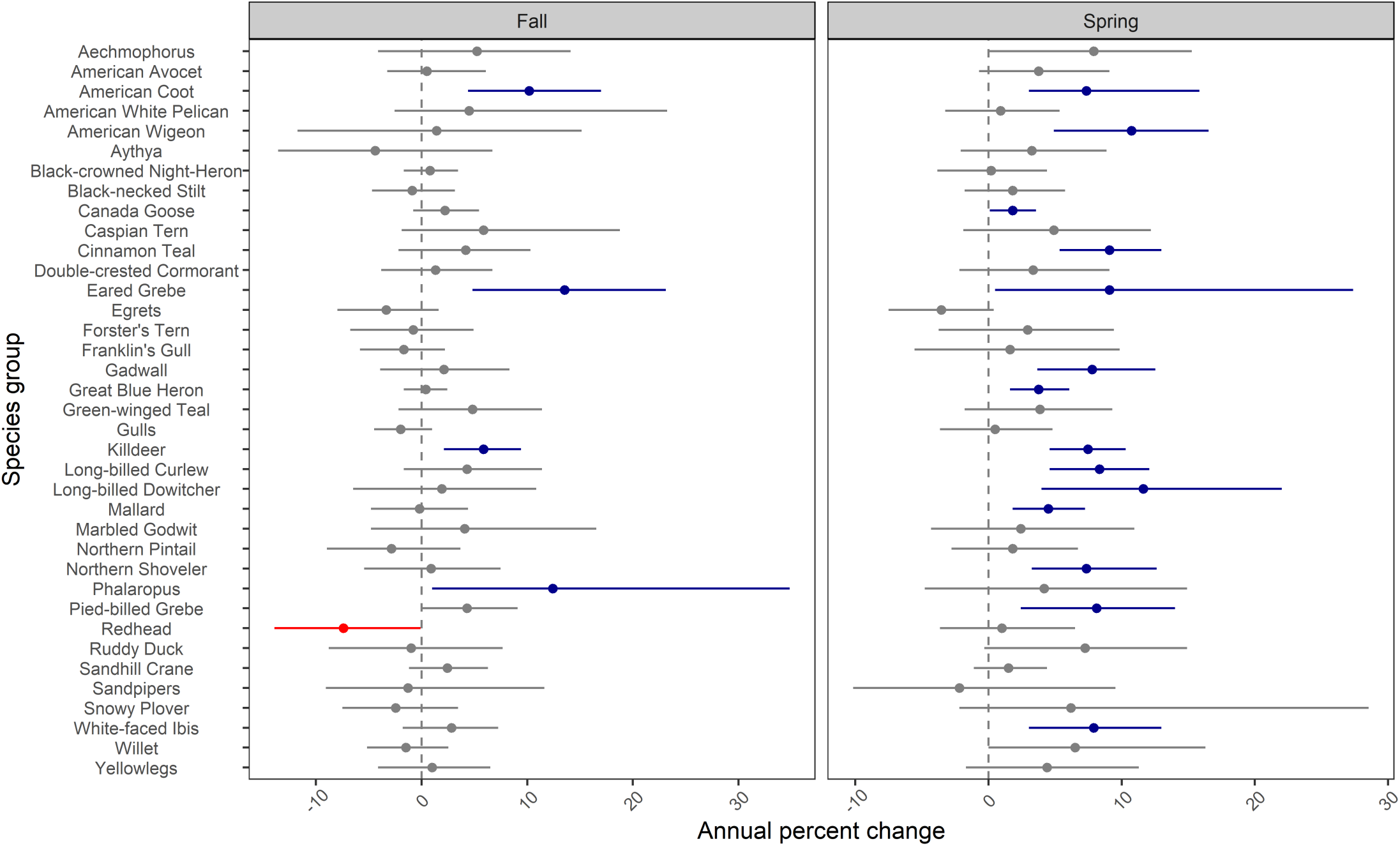
Aggregate fall and spring annual trends for shorebird, waterfowl, and other waterbird species or groups across Great Salt Lake survey units between 1997 and 2017. The aggregate trend was based on summed relative abundances modeled for each survey unit annually and regressed against year. Point estimates represent the median of the posterior distributions for aggregate trends whereas horizontal bars represent the 95% credible interval. Significant positive and negative aggregate trends are in blue and red, respectively

**Fig. 3.**
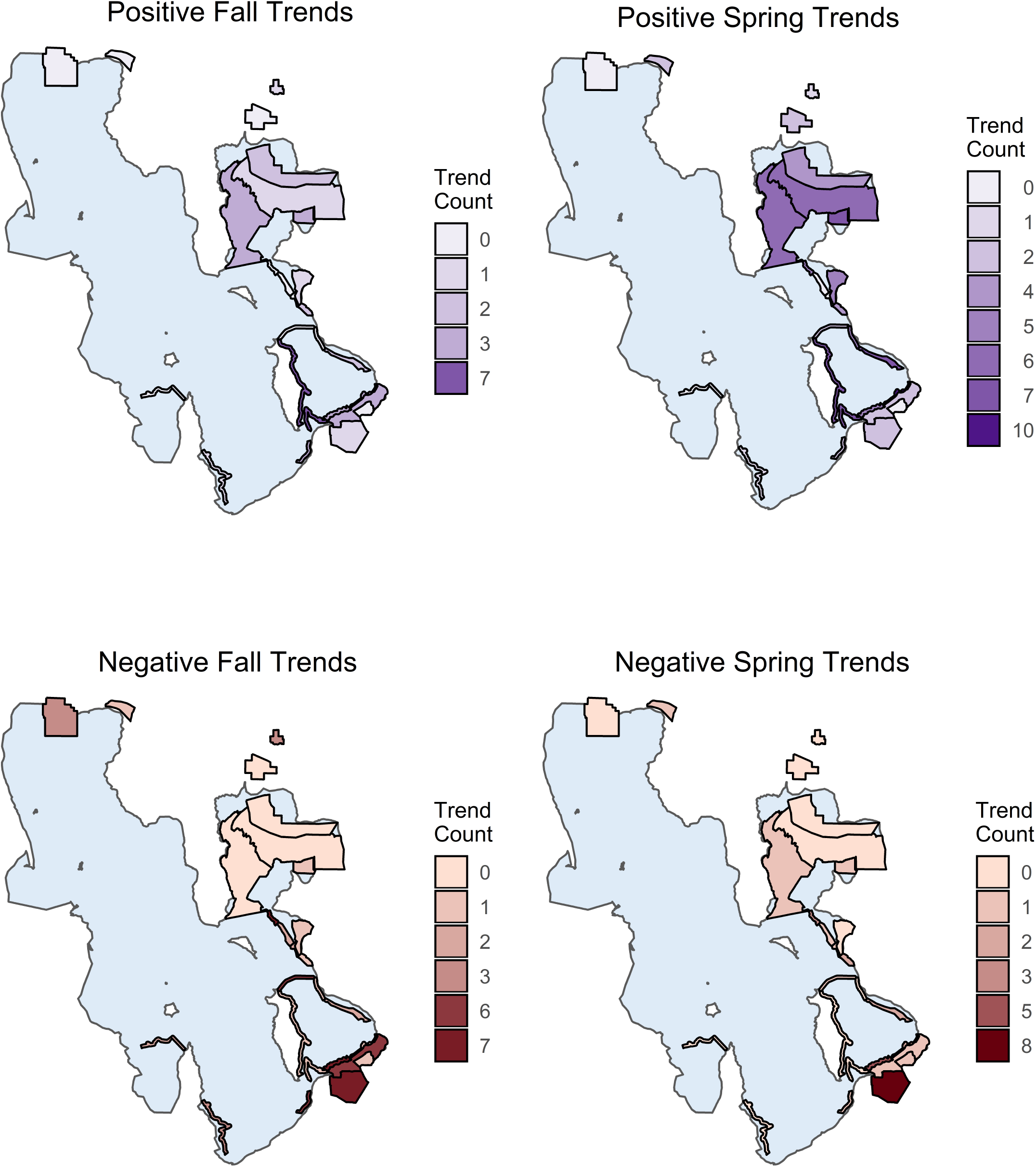
Number of positive and negative seasonal trends (1997 - 2017) for species or groups at 24 individual survey units where the Great Salt Lake Ecosystem Program conducts long-term surveys for migratory shorebirds, waterfowl, and other waterbirds at Great Salt Lake, Utah.

**Table 4.** Aggregate fall and spring annual trends for shorebird, waterfowl, and other waterbird species or groups across Great Salt Lake survey units between 1997 and 2017. The aggregate trend was based on summed relative abundances modeled for each survey unit annually and regressed against year. Significant aggregate trends are bolded, and the 95% credible interval for each trend is reported parenthetically. For each species, the numbers of survey units with significant positive (+), negative (-), or stable (S) trends are also reported. The number of survey units modeled varied between 23 and 24 depending on the species or group.

| Species or Group | Fall |  | Spring |  |
| --- | --- | --- | --- | --- |
|  | Aggregate Trend | Survey Unit (+/-/S) | Aggregate Trend | Survey Unit (+/-/S) |
| *Aechmophorus* | 5.23<br>(-4.11, 14.11) | 4/0/19 | 7.90<br>(0.00, 15.26) | 3/2/18 |
| American Avocet | 0.50<br>(-3.25, 6.08) | 0/2/22 | 3.77<br>(-0.70, 9.09) | 1/2/21 |
| American Coot | **10.19**<br>**(4.39, 17.00)** | 3/10/10 | **7.36**<br>**(3.05, 15.84)** | 4/0/19 |
| American White Pelican | 4.50<br>(-2.57, 23.24) | 2/1/20 | 0.90<br>(-3.25, 5.34) | 0/0/23 |
| American Wigeon | 1.41<br>(-11.75, 15.14) | 0/1/22 | **10.74**<br>**(4.92, 16.53)** | 7/1/15 |
| *Aythya* | -4.40<br>(-13.58, 6.72) | 0/0/23 | 3.25<br>(-2.08, 8.87) | 1/1/21 |
| Black-crowned Night-Heron | 0.80<br>(-1.69, 3.46) | 0/0/23 | 0.20<br>(-3.83, 4.39) | 1/1/21 |
| Black-necked Stilt | -0.90<br>(-4.69, 3.15) | 0/9/14 | 1.82<br>(-1.78, 5.76) | 0/0/23 |
| Canada Goose | 2.22<br>(-0.80, 5.44) | 0/0/23 | **1.82**<br>**(0.10, 3.56)** | 4/4/15 |
| Caspian Tern | 5.87<br>(-1.88, 18.77) | 1/0/22 | 4.92<br>(-1.88, 12.19) | 0/0/23 |
| Cinnamon Teal | 4.19<br>(-2.18, 10.30) | 1/1/21 | **9.09**<br>**(5.34, 12.98)** | 11/0/12 |
| Double-crested Cormorant | 1.31<br>(-3.83, 6.72) | 0/1/22 | 3.36<br>(-2.18, 9.09) | 0/0/23 |
| Eared Grebe | **13.54**<br>**(4.81, 23.12)** | 5/0/19 | **9.09**<br>**(0.50, 27.38)** | 1/4/19 |
| Egrets | -3.34<br>(-7.97, 1.61) | 0/4/19 | -3.54<br>(-7.50, 0.40) | 0/2/21 |
| Forster's Tern | -0.80<br>(-6.76, 4.92) | 0/1/22 | 2.94<br>(-3.73, 9.42) | 0/4/19 |
| Franklin's Gull | -1.69<br>(-5.82, 2.22) | 0/3/21 | 1.61<br>(-5.54, 9.86) | 0/0/24 |
| Gadwall | 2.12<br>(-3.92, 8.33) | 1/3/19 | **7.79**<br>**(3.67, 12.52)** | 8/0/15 |
| Great Blue Heron | 0.40<br>(-1.69, 2.43) | 2/0/21 | **3.77**<br>**(1.61, 6.08)** | 2/0/21 |
| Green-winged Teal | 4.81<br>(-2.18, 11.41) | 2/0/21 | 3.87<br>(-1.78, 9.31) | 4/0/19 |
| Gulls | -1.98<br>(-4.50, 1.01) | 0/3/21 | 0.50<br>(-3.63, 4.81) | 0/2/22 |
| Killdeer | **5.87**<br>**(2.12, 9.42)** | 3/0/21 | **7.47**<br>**(4.60, 10.30)** | 6/0/18 |

**Table 4 (continued).** Aggregate fall and spring annual trends for shorebird, waterfowl, and other waterbird species or groups across Great Salt Lake survey units between 1997 and 2017.
| Species or Group | Fall |  | Spring |  |
| --- | --- | --- | --- | --- |
|  | Aggregate Trend | Survey Unit (+/-/S) | Aggregate Trend | Survey Unit (+/-/S) |
| Long-billed Curlew | 4.29<br>(-1.69, 11.41) | 0/0/24 | **8.33**<br>**(4.60, 12.08)** | 7/0/17 |
| Long-billed Dowitcher | 1.92<br>(-6.48, 10.85) | 2/0/21 | **11.63**<br>**(3.98, 22.02)** | 4/0/19 |
| Mallard | -0.20<br>(-4.78, 4.39) | 4/2/17 | **4.50**<br>**(1.82, 7.25)** | 4/0/19 |
| Marbled Godwit | 4.08<br>(-4.78, 16.53) | 0/0/24 | 2.43<br>(-4.31, 10.96) | 0/0/24 |
| Northern Pintail | -2.86<br>(-8.97, 3.67) | 0/2/21 | 1.82<br>(-2.76, 6.72) | 0/0/23 |
| Northern Shoveler | 0.90<br>(-5.45, 7.47) | 1/1/21 | **7.36**<br>**(3.25, 12.64)** | 7/0/16 |
| *Phalaropus* | **12.41**<br>**(1.01, 34.85)** | 1/0/23 | 4.19<br>(-4.78, 14.91) | 0/0/24 |
| Pied-billed Grebe | 4.29<br>(-0.10, 9.09) | 2/0/21 | **8.11**<br>**(2.43, 14.00)** | 1/0/22 |
| Redhead | **-7.41**<br>**(-13.93, -0.10)** | 0/9/14 | 1.01<br>(-3.63, 6.50) | 0/0/23 |
| Ruddy Duck | -1.00<br>(-8.79, 7.68) | 0/1/22 | 7.25<br>(-0.30, 14.91) | 3/0/20 |
| Sandhill Crane | 2.43<br>(-1.19, 6.29) | 1/1/21 | 1.51<br>(-1.09, 4.39) | 0/0/23 |
| Sandpipers | -1.29<br>(-9.06, 11.63) | 0/6/18 | -2.18<br>(-10.15, 9.53) | 0/1/23 |
| Snowy Plover | -2.47<br>(-7.50, 3.46) | 0/0/24 | 6.18<br>(-2.18, 28.53) | 0/0/24 |
| White-faced Ibis | 2.84<br>(-1.78, 7.25) | 1/0/22 | **7.90**<br>**(3.05, 12.98)** | 5/1/17 |
| Willet | -1.49<br>(-5.16, 2.53) | 0/0/24 | 6.50<br>(0.00, 16.30) | 1/5/18 |
| Yellowlegs | 1.01<br>(-4.11, 6.50) | 0/1/22 | 4.39<br>(-1.69, 11.29) | 0/0/23 |

### Spring

During spring, 14 species or groups showed positive aggregate population trends from 1.82 to 11.63% (Fig. 2, Table 4). All other species or groups had stable aggregate spring trends. Twenty-one species or groups had positive local trends in up to 11 survey units, and these 21 included all 14 species or groups showing positive aggregate trends (Fig. 2, Table 4). The unit with the largest number of positive local spring trends for species and groups was located in Farmington Bay (Fig. 3). Thirteen species or groups displayed negative local spring trends in as many as 5 survey units, and of these 13, four possessed positive aggregate trends (Table 4). The greatest number of negative local spring trends for species and groups occupied the southernmost survey unit associated with Farmington Bay (Fig. 3). Foraging technique (χ^2^_2,27_ = 3.25, p = 0.24), migration strategy (χ^2^_2,30_ = 2.33, p = 0.15) and taxonomic group (χ^2^_2,30_ = 1.10, p = 0.72) were unassociated with aggregate trend direction.

## Discussion

Great Salt Lake and its associated wetlands are recognized for their importance to shorebirds, waterfowl, and other waterbirds (Aldrich and Paul 2002; Chipley *et al*. 2003; Wilsey *et al*. 2017), but relatively few studies have examined long-term trends for bird species at the lake (Sloan 1982; Paton 1997; King and Anderson 2005; Neill *et al*. 2017). Using a 21-year dataset (1997 – 2017), we examined trends in high-use and presumably, relatively high habitat quality areas of Great Salt Lake and its associated wetlands, and accordingly, we predicted a predominance of stable trends (Gill *et al*. 2001). We found that 36 of 37 bird species or species groups displayed stable or positive trends during fall and spring at Great Salt Lake. These stable and positive trends, in conjunction with potential declines of some species at other saline lakes experiencing declining water levels (Larson *et al*. 2016; Jones *et al*. 2019), further emphasize the importance of these survey units in or associated with Great Salt Lake to migratory, water-dependent birds.

Intentional efforts to maintain stable and positive trends will require identifying factors regulating populations, designing and implementing conservation and management actions to prevent factors from limiting populations, and monitoring the success of actions. Given that the birds examined are migratory, ecological and demographic factors contributing to Great Salt Lake counts and trends might originate in other areas and seasons (Newton 2004). While many factors can affect trends at Great Salt Lake and other saline lakes, one receiving considerable regional and local attention is a reduction of water inflows affecting habitat amount, timing, and quality (e.g., by changing salinity) (Ivey and Herziger 2006; Wilsey *et al*. 2017; Senner *et al*. 2018; Haig *et al*. 2019). Securing adequate water supplies at key times of year is paramount to provide and sustain local and regional habitats for shorebirds, waterfowl, and other waterbirds (Ivey and Herziger 2006; Wilsey *et al*. 2017).

Stable and positive aggregate trends were not directly transferable to individual survey units as some species with stable or positive aggregate trends displayed negative trends in individual survey units. As an example, the Black-necked Stilt (*Himantopus mexicanus*) a shorebird species of regional focus (Thomas *et al*. 2013), declined significantly in nine survey units during fall despite showing a stable aggregate trend (Table 4). There were survey units in Farmington and Ogden bays that possessed relatively large numbers of declining trends for species or species groups, such as Franklin’s Gull (*Leucophaeus pipixcan*) and Willet (*Tringa semipalmata*). Such observations based on trends for individual survey units can inform prioritizations of species and areas for conservation and management. Further, the presence of stable, positive, and negative trends across individual survey units offers an opportunity to evaluate environmental drivers of changing species and group counts, and there are many candidates to consider, including the composition and dynamics of vegetation (Rohal *et al*. 2018), hydrology (Cavitt 2013), food resources (Conover and Caudell 2009), and others (Ma *et al*. 2010). Relationships between trends and environmental drivers could assist with the development of conservation or management strategies and tactics to maintain aggregate stable or positive trends.

When established, the present bird monitoring program was designed to permit trends to be examined, but evaluations of correlations with environmental drivers at the scale of individual survey units were not within scope (Paul and Manning 2002). Consequently, data on potential drivers are not currently collected along with bird counts. There are long-term environmental monitoring efforts that temporally coincide with bird surveys, such as river inflow and lake elevation monitored by the U.S. Geological Survey (U.S. Geological Survey 2020), and previous studies have demonstrated the potential of relating these environmental data to bird counts for specific areas of Great Salt Lake (Cavitt 2013). Another potential source of environmental data is the use of remotely sensed imagery to map features of interest, such as habitat covers and changing vegetation composition (Long *et al*. 2017). Future efforts are needed to evaluate the feasibility of concurrently monitoring birds and environmental drivers, linking bird counts and temporally coincident environmental datasets, and possibly developing needed environmental datasets through remote sensing datasets and techniques. The authors are currently analyzing linkages among species and group counts and hydrological variables.

We did not find evidence that taxonomic group, migration strategy, or foraging technique affected the odds of a species displaying a positive versus stable trend. At times, our taxonomic groupings are used as the basis for local and regional management planning (Dybala *et al*. 2017; Tavernia *et al*. 2017; LMVJV Shorebird Working Group 2019), and foraging characteristics have been suggested or used to group birds to evaluate responses to management actions or environmental perturbations (Verner 1984; De Graaf *et al*. 1985; DeLuca *et al*. 2004). Given the greater distances they cover on an annual basis, longdistance migrants might be more sensitive than are short-distance migrants to environmental disturbances caused by climate change (cf. Galbraith *et al*. 2014). Species within taxonomic groups, migration strategies, or foraging groups can differ in specific habitat requirements, demography, geography, and the spatial and temporal scales at which they respond to their environments (Block *et al*. 1987; Noon *et al*. 2009), so species within these groups should not necessarily be assumed to show similar trends. Given the different species-specific trend directions within our groupings, we cannot recommend our taxonomic, migration strategy, or foraging technique groupings as meaningful for conservation or management planning at Great Salt Lake and its associated wetlands. While our specific migration strategy classification did not relate to trend direction, we support calls for coordinated research, monitoring, and management across multiple sites and seasons to address the needs of migratory birds throughout their annual cycles (Runge *et al*. 2014; Marra *et al*. 2015).

For our trends analysis, we assumed that counts represent a constant proportion of the individuals within a survey unit across years. The proportion counted is a function of detection probability, or the probability of correctly identifying an individual present within the survey unit (Thompson 2002). If our assumption holds, our trends represent changes in numbers of individuals using survey units rather than being an artefact of a systematic change in detection probability. A variety of factors affect detection probabilities including species identity, observer identity and experience, survey method, habitat conditions (e.g., percentage of open water), season, time of day, weather conditions, ambient noise, and others (Johnson 2008). We do not attempt count comparisons between species, and thus, we are unconcerned about likely differences in detection probabilities across species. To the extent possible, we have attempted to control other factors through personnel decisions and standardized protocols.

The professional status of observers (i.e., wildlife biologists versus volunteers) has remained constant for the majority of survey units whereas individual observers may turnover from year-to-year. Personnel with long-term monitoring experience supervise counts of their respective survey units each year. Thus, we expected turnover in observers to increase the variability of counts, but we did not expect a consistent, directional change in detection probability because of changing observers. The implementation of an intensive training program for observers would reduce identification errors and variability among observers (Kepler and Scott 1981; Greenwood 2007). Training could help with identification of species that are currently lumped into groups. Survey methods and travel modes have changed for some units to different degrees with time, especially following 2001 surveys. However, including a random factor to control for pre-2001 and post-2001 survey status did not significantly change the proportions of positive or stable trends observed (unpublished data). Our survey protocols detail the seasonal and daily timing of surveys, and there has been no systematic change in survey timing from 1997 through 2017. An increased frequency of surveys during spring and fall seasons would reduce variability in counts across years. For individual species, our survey periods may be aligned to greater or lesser degrees with their primary periods of migration, and thus, where possible, we recommend comparing our trends to trends from species-specific survey efforts (e.g., for fall migration of the Eared Grebe, Neill *et al*. 2017). Acceptable weather conditions are specified with surveys not occurring if winds exceed a Beaufort scale of three (18.5 km/hr) or precipitation is more than intermittent rain. While ambient noise levels are not recorded, the majority of species and individuals are observed visually, and for that reason, we do not expect potential increases in ambient noise to have biased outcomes from trend analyses. Thus, unless an unknown and uncontrolled factor is acting to affect detection probability, our trends reflect changes in the numbers of individuals using survey units.

We caution against an uncritical interpretation of the stable and increasing bird population trends at Great Salt Lake. Playas, an important shorebird habitat, are not currently represented among the established survey units and are threatened by human development (Sorensen *et al*. 2020). Further, within existing survey units, stable or increasing counts for a species within a fixed area (i.e., density) are not necessarily indicators that habitat quality is stable or improving (Van Horne 1983). As examples, birds may elect to settle in lower quality habitat as preferred habitat becomes saturated with conspecifics (Gill *et al*. 2001), or density may increase in remaining areas as habitat is lost from a landscape (Hagan *et al*. 1996). Metrics of habitat quality should assess both density and demographic measures of individual performance, i.e., per capita survival and reproduction, within an area (Van Horne 1983).

During migration, refueling rates and the amount of fuel deposited at stopover locations influence how long a bird must remain at a stopover and may affect subsequent survival and reproduction during the remainder of its annual cycle (Drent *et al*. 2003; Baker *et al*. 2004; Newton 2006). Accordingly, regional management planning for migrating waterfowl and shorebirds often uses bioenergetics models to quantify the amount of foraging habitat and food resources required to meet explicit population objectives (e.g., Petrie *et al*. 2013; Dybala *et al*. 2017; LMVJV Shorebird Working Group 2019). Petrie *et al*. (2013) applied a bioenergetics model to assess the adequacy of foraging habitat and food resources for nonbreeding waterfowl at Great Salt Lake. They found that food supply adequacy varied across waterfowl guilds, managed versus unmanaged lands, periods and seasons, and projected hydrologic conditions. Petrie *et al*. (2013) indicated that model improvements could be achieved with better estimates of wetland productivity and data on waterfowl resource selection. Despite interest in a bioenergetics model for shorebirds at Great Salt Lake (Thomas *et al*. 2013), no such efforts have been completed to date. To our knowledge, the bioenergetics needs of only a single waterbird species, the Eared Grebe (Conover and Caudell 2009), have been assessed in detail. We support applying and improving bioenergetics approaches at Great Salt Lake to identify clear habitat benchmarks to meet the foraging needs of migrating individuals and, consequently, to prevent foraging conditions at the lake from limiting bird populations.

Our trends analyses indicate that 36 of 37 examined species or species groups were stable or increasing during a 21-year period in the surveyed areas at Great Salt Lake, a site identified as important for shorebirds, waterfowl, and other waterbirds (North American Waterfowl Management Plan Committee 2004; Ivey and Herziger 2006; Petrie *et al*. 2013). Maintaining current trends will require setting explicit population (e.g., trend-based) objectives, identifying environmental factors potentially limiting populations, designing and implementing actions to address limiting factors, monitoring action outcomes, and adapting objectives, actions, and monitoring as needed based on observed outcomes (National Ecological Assessment Team 2006). Monitoring might be expanded to other low bird-use areas of the lake to facilitate detecting early species declines before they manifest themselves in highly used areas of presumably high habitat quality. Adding survey units in unrepresented habitat types (e.g., playas), additional duck clubs, and inland wetland areas would provide a more complete picture of wetland-dependent bird status and trends at Great Salt Lake. Careful consideration should be given to relationships among water inflows, lake elevation, and the ability of Great Salt Lake and its associated wetlands to meet the energetic needs of migratory shorebirds, waterfowl, and other waterbirds. Ultimately, the needs of migratory birds must be met throughout their annual cycles, so management and monitoring efforts at Great Salt Lake must be nested within larger regional and continental plans and programs to address shorebirds, waterbirds, waterfowl, and their habitats.

## Acknowledgments

We thank all the biologists, field technicians, and volunteers for collecting the field data used in these analyses. A. Manning, D. Paul, and C. Perschon provided key leadership in the early years of the monitoring program. P. Birdsey, G. Evans, F. Howe, K. Johnson, D. Mann, R. Norvell, and K. Poulsen helped with survey and database design and management. We thank A. Neville, C. Wilsey, E. Sorensen, K. Stockdale, M. Malmquist, M. Shoop, and S. Senner for comments that improved the manuscript.

## Notes

### Competing Interest Statement

The authors have declared no competing interest.

